# Design of Tau Aggregation Inhibitors Using Iterative Machine Learning and a Polymorph-Specific Brain-Seeded Fibril Amplification Assay

**DOI:** 10.1101/2025.03.09.642189

**Authors:** Alessia Santambrogio, Robert I. Horne, Michael A. Metrick, Z. Faidon Brotzakis, Dillon Rinauro, Ergina Vourkou, Katerina Papanikolopoulou, Efthimios M. C. Skoulakis, Sara Linse, Byron Caughey, Michele Vendruscolo

**Author notes:** Authors contributed equally to this manuscript (Alessia Santambrogio, Robert I. Horne, Michael A. Metrick).

## Abstract

The aggregation of tau into amyloid fibrils is associated with Alzheimer’s disease (AD) and related tauopathies. Since different tauopathies are characterised by the formation of distinct tau fibril morphologies, it is important to combine the search of tau aggregation inhibitors with the development of *in vitro* tau aggregation assays that recapitulate aggregation as it may occur in the brain. Here we address this problem by reporting an *in vitro* tau aggregation assay in which AD brain homogenates are used to seed the generation of first-generation tau fibrils in a polymorph-specific manner under quiescent conditions. These fibrils are then used to create amyloid seed libraries from which second-generation kinetic assays can be readily performed. Using this strategy, we illustrate an iterative machine learning method for the identification of small molecules for the polymorph-specific inhibition of the *in vitro* formation of tau fibrils. We further show that the small molecules selected by this procedure are potent inhibitors in a *Drosophila* tauopathy model.

## Introduction

Alzheimer’s disease (AD) is characterised by the presence of amyloid deposits that spread across the brain in specific spatio-temporal patterns (*1-7*). Preventing such propagation is a common drug discovery strategy for the treatment and prevention of AD and related neurodegenerative disease (*8-10*).

A promising therapeutic approach is based on the observation that a fundamental property of amyloid aggregates is their ability to promote the formation of new aggregates (*11, 12*). Because this autocatalytic process is dependent on the presence of amyloid fibrils, their structures are likely to determine its efficiency (*13-15*). However, through cryo-electron microscopic (cryo-EM) studies, it is now known that tau can adopt over 20 different polymorphic amyloid structures throughout the various tauopathies (*16-18*). Since such polymorphism creates significant challenges for drug discovery, it is important to develop *in vitro* assays of tau aggregation that result in disease-specific fibril polymorphs (*19*). This is difficult because, depending on the conditions (pH, salt, temperature, cofactors) and the sequences of the tau isoforms, *in vitro* assays might generate fibril structures that depend on the respective solution conditions, many of which result in amyloid fibrils of morphologies not observed in brain extracts of tauopathy patients (*19-21*)

To reproduce the amyloid structures found *in vivo*, a possible approach is to develop assays whereby brain-derived seeds are used to induce recombinant monomeric tau to adopt fibrils of specific morphologies, either *in vitro (22, 23*) or in cell systems (*24*). This strategy relies on the observation that the free energy barrier for elongation is lower than that for the spontaneous generation of new seeds (*11, 25*), and it should therefore be possible to propagate a conformer through seeding, even under conditions where another more stable conformer may arise from primary nucleation (*26, 27*). This method was recently adopted to obtain α-synuclein fibrils with the morphology observed in multiple system atrophy (MSA) (*23*), which is distinct from those observed in Parkinson’s disease (PD) or dementia with Lewy bodies (DLB) (*28*). However, although one protofilament of mature fibrils was faithfully propagated, the corresponding counter-filaments in many cases adopted structures not observed in reconstructions of MSA fibrils from patient brain extracts as elucidated via cryo-EM (*29*).

An alternative method is to identify solution conditions for *in vitro* aggregation assays that lead to the recreation of the amyloid fibril morphologies observed in disease through spontaneous seed formation, in the absence of brain-derived seeds. Significant progress has been recently made in this direction with tau under shaking conditions (*19*). For the application of this approach to drug design, however, the *in vitro* assay should ideally also be consistent with the conditions *in vivo*, and thus be conducted without vigorous shaking, relying only on gentle agitation brough about by moving the samples in the fluorescence reader (*30*) to identify candidate inhibitors with a clinically-relevant mechanism of action.

Here we report a framework for addressing this problem and its application to drug design. This framework is based on a careful kinetic analysis (*31, 32*) under quiescent reaction conditions whereby individual microscopic processes might be specifically targeted with small molecule inhibitors (*25, 33, 34*). In the first step, we identified small molecule candidates with an *in silico* docking procedure to a hydrophobic pocket of a previously reported tau fibril structure from AD patients (*16*). This pocket spans a sequence next to a region known to be critical to *in vitro* paired-helical filament propagation (PHF6, VQIVYK) (*18*). In the second step, we used a kinetic analysis of tau aggregation (*25*) coupled with iterative machine learning (*35*) to enhance the potency of the candidate inhibitors. In the third step, we confirmed the conformational specificity by observing no inhibition of Pick’s disease (PiD)-seeded tau aggregation, and we verified the *in vivo* potential of the aggregation inhibitors by rescuing viability in a *Drosophila* model of tauopathy (*36, 37*). Taken together, our results illustrate the use of machine learning methods to identify compounds capable of targeting disease-specific polymorphs in tauopathies.

## Results

### Brain-derived polymorph-specific tau seed amplification and biophysical characterisation of the resulting aggregates

In this work, we used the approach illustrated in **Figure S1** to achieve the *in vitro* amplification of polymorph-specific tau seeds. In a first amplification step (producing first-generation recombinant tau fibrils), we used brain-derived seeds and monomers of K12, a 3-repeat (3R) fragment of tau previously designed to amplify tau aggregates from crude AD and PiD brain homogenates (*26*). The method was adapted to avoid the use of heparin, as this cofactor was shown to lead to the formation of structures different from those observed in diseases (*20*). We incubated 32 AD, 32 PiD and 32 cerebrovascular vascular disease (CVD) brain homogenate-seeded reactions (0.0001% w/v) with rounds of 500 rpm shaking and rest in 250 mM trisodium citrate and 10 mM HEPES at pH 7.4 (**Figure 1A**). Characteristic differences in thioflavin T (ThT) amplitudes between AD-seeded and PiD-seeded reaction products were observed, as previously reported (**Figure 1B**) and kinetic acceleration of lag times ahead of control reactions seeded with CVD was achieved (**Figure 1C**). We sought further biophysical characterisation of the K12 aggregates with ATR-FTIR analysis. We previously reported that differences in ThT fluorescence intensities between products of AD-seeded and PiD-seeded reactions correlated with secondary structural differences visualised by ATR-FTIR analysis (*26*). Characteristic 1630 cm^-1^ and 1618 cm^-1^ β-sheet vibrational modes were observed for AD-derived fibrils, which were distinct from PiD reaction products with prominent 1633 cm^-1^ and 1628 cm^-1^ vibrations (**Figure 1D**). The amplification of PiD-seeded K12 reactions alongside those of AD served as a control to enhance the robustness of divergent conformer amplification.

**Figure 1.**
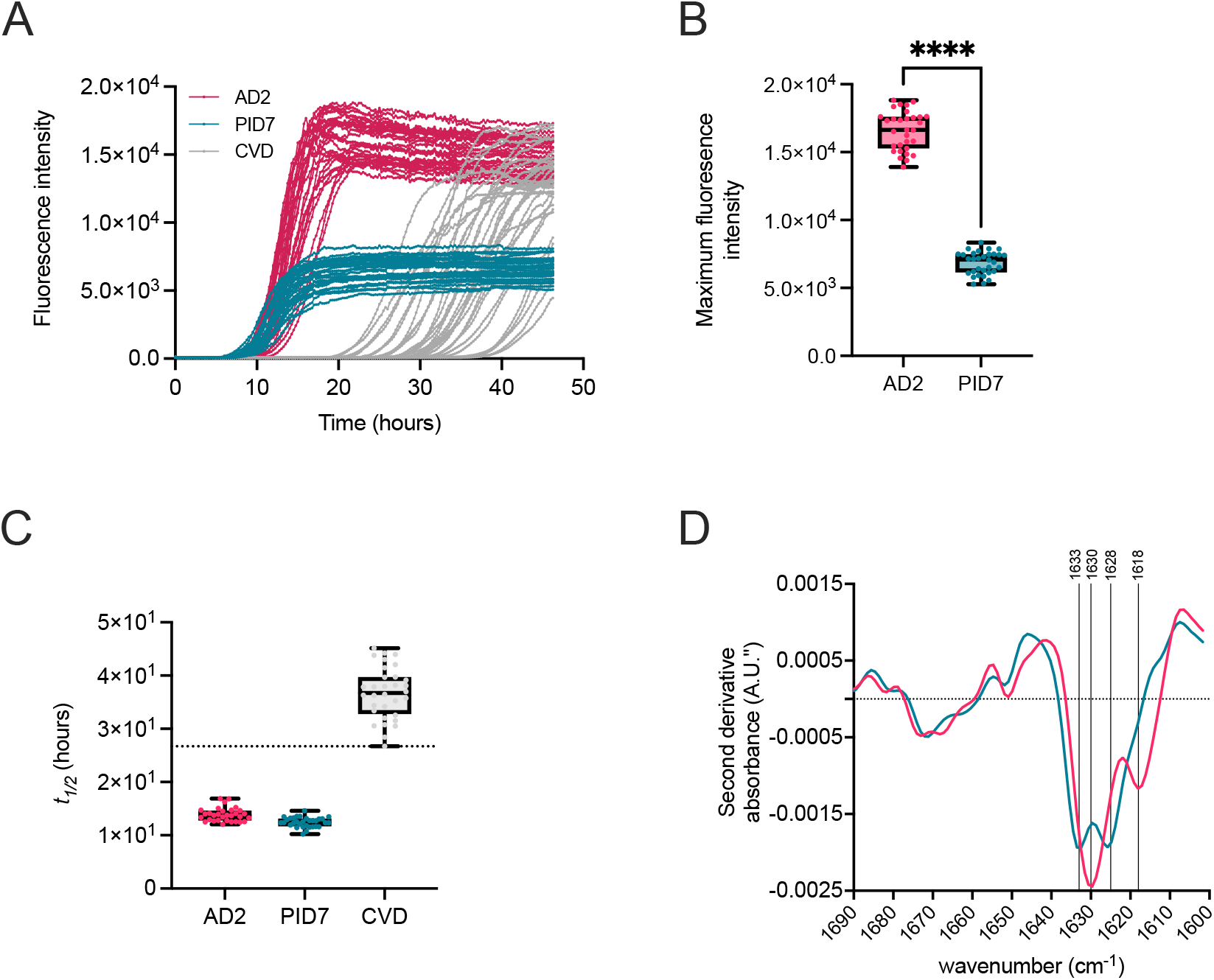
Biophysical characterization of first-generation K12 tau fibrils. (**A**) Fibrils were generated by seeding conversion of recombinant K12 tau with AD brain homogenates (pink), PiD brain homogenates (teal) or CVD brain homogenates (grey). (**B**) ThT fluorescence maxima of amplified products. Differences of means were analysed by a one-way ANOVA test. (**C**) Boxplots of the half-time (*t*_*1/2*_) of the reactions in panel A. (**D**) Second derivative ATR-FTIR spectra of pooled reactions with raw traces and ThT maxima corresponding to panels A and B.

### Kinetic analysis of polymorph-specific tau propagation

We modelled the aggregation process using a global fitting method (*32*) to identify targetable microscopic mechanisms in the propagation of AD-derived tau aggregates. First-generation AD-derived fibrils were used as seeds in a dilution series of monomeric K12 to study such pathways under quiescent conditions (**Figure 2A,B**). The monomer dependence of half times of aggregation is shown in **Figure 2C**. Kinetic mechanisms not consistent with the observed data were removed from consideration (*32*). The slope (γ) of the double-logarithmic plot of monomer concentration versus half time approached the value of −0.5, suggesting that in the AD-seeded aggregation assay tau aggregation involves secondary processes such as secondary nucleation and fragmentation, in addition to elongation (*25, 32*). Fibrils were recovered from the quiescent reactions and visualized by transmission electron microscopy (TEM) (**Figure 2D**).

**Figure 2.**
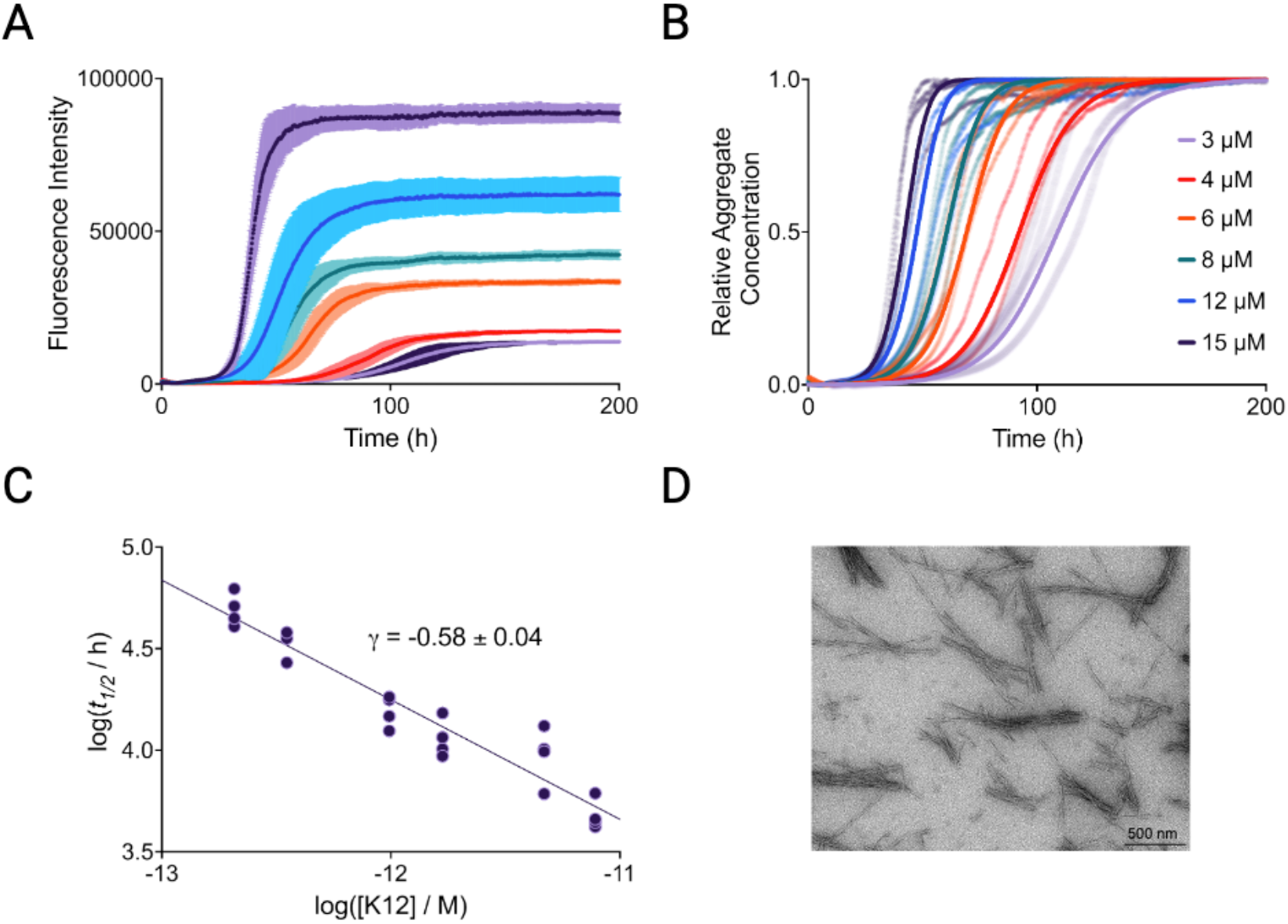
Kinetic analysis of polymorph-specific tau propagation. (**A**) Kinetic traces for AD-derived fibrils seeded into a dilution series of monomeric K12 at 3 μM (lilac), 4 μM (red), 6 μM (orange), 8 μM (teal), 12 μM (blue) and 15 μM (purple). Error bars indicate the SD. (**B**) Normalised kinetic traces (points) with overlaid fits (solid lines) of a secondary process-dominant aggregation mechanism. (**C**) Scaling of the half-time of the aggregation reaction with the concentration of K12. The scaling exponent, γ, had a value close to −0.5, implying a low dependence on monomer concentration, and dominating secondary processes. (**D**) TEM images of AD brain homogenate-seeded K12 fibrils grown quiescently.

### Prediction of small molecule inhibitors from *in silico* docking on AD tau fibrils

Candidate small molecules were identified via docking to cryo-EM structures (PDB 5O3L) of tau fibrils purified from brain extracts of patients with AD(*16*). Our hypothesis was that molecules that bind to the fibril structures could modulate aggregation processes involving formation of new misfolded aggregates from existing ones (*35, 38, 39*). Potential binding pockets were identified via the pocket identification software Fpocket (*40*), and pockets of low solubility were prioritised with CamSol (*41*), identifying areas which would be likely to promote protein-protein interactions between monomer and fibrils (see Methods and **Figure S2**). **Figure S3A** highlights the surface-exposed hydrophobic region adjacent to the PHF6 region of tau (*18*) which we chose for docking small molecules. A subset of ∼1.5 million small molecules from the ZINC (*42*) database was selected which passed CNS MPO criteria (*43*), a metric of the propensity of a small molecule to reach the central nervous system. The predicted binding affinity of this subset to the chosen binding site was calculated via AutoDock Vina (*44*) docking software The top 10,000 docked small molecules from this screening were then redocked using FRED (*45*) (OpenEye Scientific Software) to improve confidence by consensus scoring of the top hits, and the best predicted binders (∼120) were obtained for experimental testing.

### Polymorph-specific inhibition of aggregation processes using the predicted small molecule binders

We investigated the effects of the predicted small molecule binders on the microscopic processes of tau aggregation. Aggregation kinetic experiments were conducted at 5 μM K12 monomer concentration in the presence of 50 nM (monomer equivalents) first-generation tau fibril seeds, with addition of ± 20 μM small molecule. The primary metric of potency in the aggregation assays was the normalised half time (*t*_1/2_) of aggregation, the point at which half of the monomer present at the start of the aggregation has converted to amyloid fibril in the presence of an inhibitor, normalised to the negative control (DMSO) *t*_1/2_. Larger *t*_1/2_ values therefore represent aggregation inhibitors, while smaller *t*_1/2_ values represent inducers. The *t*_1/2_ values for the docked compounds (**Figure S4A**) and iterative rounds of kinetic-informed machine learning (**Figure S4B-D**) show that the iterative process progressively improves the potency of the compounds. Green bars represent potent aggregation inhibitors (*t*_1/2_ > 1.5, defined as a hit), and purple bars represent highly potent aggregation inhibitors (*t*_1/2_ > 2). Of the 105 small molecules experimentally tested in K12 AD aggregation assays, two were hits (∼2% hit rate).

These 2 hits (labelled d0 and d1) were taken forward for optimisation via a machine learning pipeline (*35*). For this stage, we defined the optimization rate in the same way as the hit rate, with any molecules exceeding *t*_1/2_ > 1.5 informing subsequent rounds of machine learning, as previously described(*35*). Of the further 32 small molecules tested in iteration #1, one was potent (3%), and another one highly-potent (3%). In iteration #2, we tested 45 small molecules, of which were 11 potent (24%), and 3 highly-potent (7%), In iteration #3, we tested 59 small molecules, yielding 10 potent (17%) and 6 highly-potent (10%) small molecules. We note that the potency of the top small molecules increased with each iteration (**Figure 3**).

**Figure 3.**
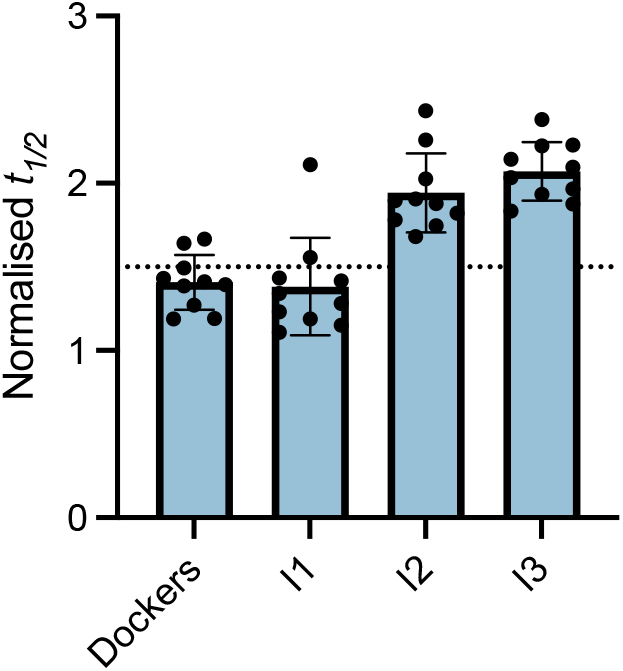
Results of the docking and iterations of the machine learning drug discovery strategy. Normalised half-time (*t*_1/2_) of AD-seeded K12 reactions for the top 10 potent aggregation inhibitors at 20 µM from different stages: docking, iteration 1, iteration 2 and iteration 3. The horizontal dotted line indicates the boundary for potent lead classification, which was normalized *t*_1/2_ = 1.5. For the docking, 105 molecules were tested, while for iterations 1, 2 and 3, the number of molecules tested was 32, 45 and 49, respectively.

Further experiments were carried out after initial screening to validate the mechanism of inhibition with the most promising hits from iteration 1 (**Figure 4A**). By modifying the input seed:monomer concentration ratio, experiments were designed to quantify perturbations in secondary processes and heterologous nucleation (low-seed experiments, **Figure 4B**) and elongation rates (high-seed experiments, **Figure 4C**). Based on their high *t*_1/2_ values, three compounds were chosen for detailed kinetic analysis: I1.21, I1.51 and I1.114 (whose chemical structures are shown in **Figure 4A**). These share high similarity (being in the same cluster upon performing Tanimoto clustering with cutoff 0.78) as they exhibit a common substructure (**Figure S3E**). The docked poses against the AD fibril structure at the binding site V313-L315 are shown in **Figure S3B-D**. All three compounds exhibited significant dose-dependent inhibition of aggregation in the low seeded experiments (**Figure 4B**), while I1.21 also exhibited dose-dependent inhibition of elongation (**Figure 4C**). When plotted and fitted to a secondary nucleation model, I1.21, I1.51 and I1.114 reduced secondary nucleation rate constant (k_2_) by 97%, 94% and 95%, respectively (**Figure S5A**) at 20 μM (4:1 stoichiometry). Additionally, I1.21 decreases the elongation rate constant (k_+_) by 29%, compared to reductions of 12% and 6% for I1.51 and I1.114, respectively (**Figure S5B**). Given that secondary processes may be the dominant mechanism in the production of misfolded oligomers associated with disease pathology, reduction of k_2_ would be desirable in potential therapeutics.

**Figure 4.**
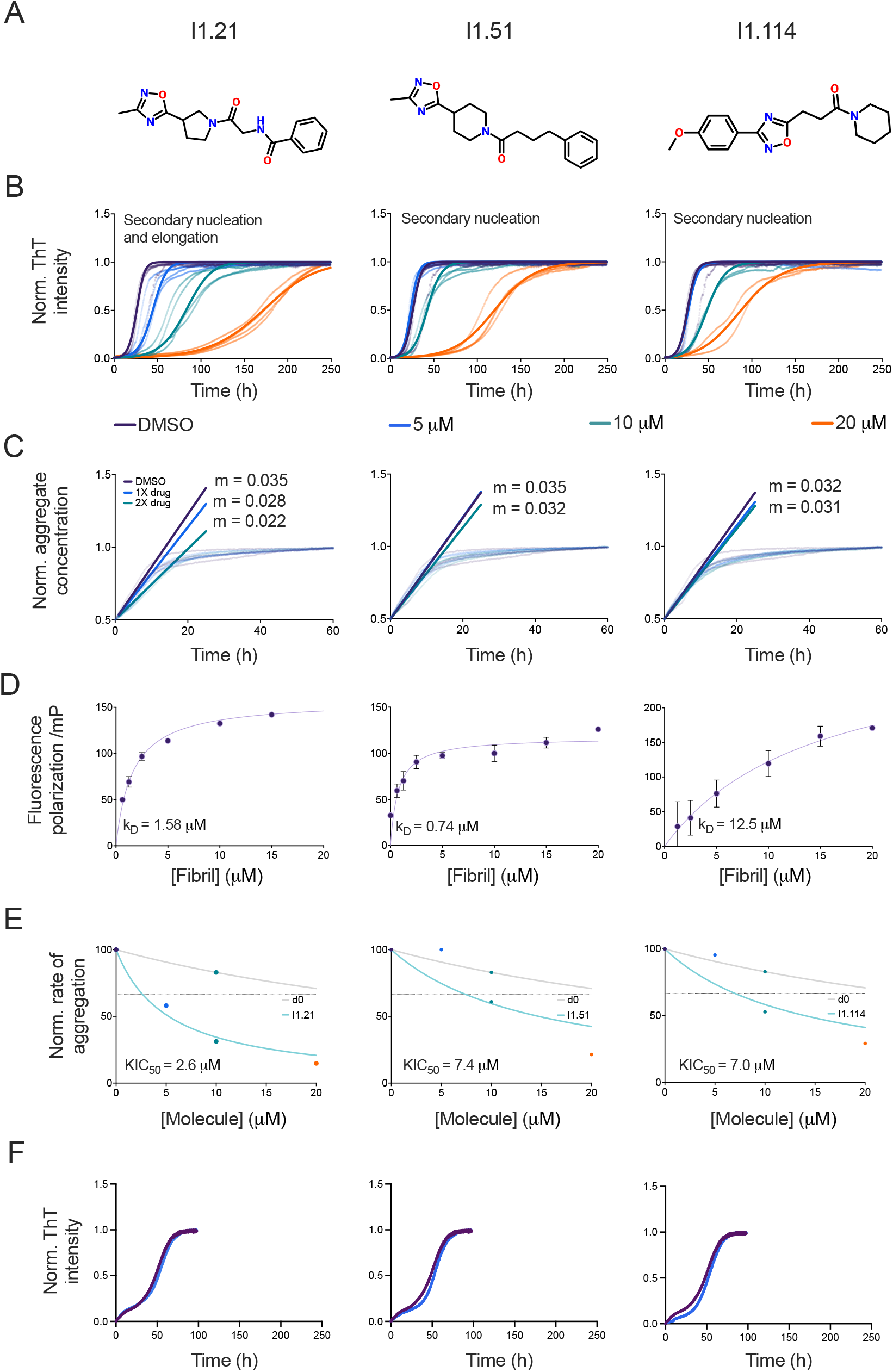
Kinetic analysis of three polymorph-specific tau aggregation inhibitors identified through iterative machine learning. (**A**) Chemical structures of compounds I1.2, I1.51 and I1.114. (**B**) Quiescent aggregation of 5 μM K12 tau seeded with 50 nM first-generation K12 tau seeds from AD-brain seeded reactions (dotted curves) in the presence of 1% DMSO (purple points), and increasing concentrations (coloured points) of either I1.21, I1.51 or I1.114; solid lines represent a multistep secondary nucleation kinetic analysis of polymorph-specific tau aggregation (*32*). A fragmentation model (*32*) also provides a similar degree of fitting. The elongation rate (*k*_*+*_) for these models was derived from (**C**) showing kinetic traces for 2.5 μM seed, 5 μM monomer AD aggregate K12 reactions in the presence of 1% DMSO (purple points), and either I1.21, I1.51 or I1.114 (coloured points). Simple linear regression analysis (solid curves) of the first 10 h of reactions in (**C**) reflect drug effects on seeded K12 elongation rates. (**D**) Change in fluorescence polarisation (in mP units) of 10 µM I1.21, I1.51 or I1.114 with increasing concentrations of K12 fibrils (concentrations given in monomer equivalents). Error bars indicate the SD. The solid line is a fit to the points using a one-step binding curve, estimating a K_D_ of 1.58 ± 0.15 μM (SD) for I1.21, 0.74 μM ± 281 nM (SD) for I1.51 and 12.5 μM ± 9.6 μM (SD) for I1.114. (**E**) Approximate rate of reaction (taken as 1/*t*_*1/2*_, normalised between 0 and 100) in the presence of 2 different molecules, the original docking hit d0 (grey), and either I1.21, I1.51 or I1.114 derived from it (light blue). The KIC_50_ of I1.21 (2.6 μM), I1.51 (7.4 μM) or I1.114 (7.0 μM) are indicated by the intersection of the fit and the horizontal dotted line. (**F**) Quiescent aggregation of 5 μM K12 tau seeded with 50 nM first-generation K12 tau seeds from PiD-brain seeded reactions and molecules at 20 µM versus 1% DMSO alone (dark purple).

The binding affinity of the three compounds was tested by fluorescence polarisation in the presence of first-generation K12 seeds over increasing fibril concentrations (**Figure 4D**). I1.51 exhibited the strongest binding, K_D_ = 0.74 μM, followed by I1.21 K_D_ = 1.58 μM, and I1.114 K_D_ = 12.5 μM. All three compounds also exhibited up to 10-fold improvement in potency over their parent compounds as shown by the rate constant plots and corresponding kinetic inhibitory concentration (KIC_50_) values (*33*) (**Figure 4E**).

To test the specificity of the three candidate compounds, we performed seeding assays of PiD-seeded K12 tau (**Figures 4F** and **S7**) and Aß42 (**Figure S8**), another misfolding-prone protein implicated in the progression of AD (*3*). We observed no significant inhibition of aggregation in both assays, confirming the specificity of the candidate compounds

### Effects of the compounds in a *Drosophila melanogaster* tauopathy model

We evaluated the effects of the compounds in a well-established *Drosophila melanogaster* tauopathy model expressing human tau 0N4R (see Methods). Measuring the viability ratio and performance index in flies treated with the leading three compounds, we found a dose-dependent reversal of tau-mediated toxicity and dysfunction (**Figures 5** and **S9**). The primary read-out for efficacy was the viability, scored as the number of flies reaching adulthood. As previously reported (*46*) pan-neuronal overexpression of tau resulted in 50% lethality (**Figure 5A**, elav/0N4R vs elav/+ p=1×10^−7^, n≥8). Treatment with 10 nM and 100 nM I1.21 restored viability to non-transgenic control levels (p=1 in both cases, n=7), while at 1 nM and 250 nM concentrations this compound had no effect (**Figure 5A**, p=2×10^−9^ and p=1×10^−5^ versus elav/+, respectively, n= 7). The lack of efficacy at 250 nM may be attributed to dose-saturation effects or altered pharmacokinetics, where higher concentrations fail to proportionally increase the therapeutic effect of the compound and may even reduce efficacy due to off-target binding. Treatment with I1.51 failed to restore viability at any of the tested concentrations (**Figure S9Ai**), whereas I1.114 restored viability to levels comparable to the non-transgenic control (elav/+) at 1 nM, 10nM and 100 nM (**Figure S9Bi**). Detailed statistics can be found in **Table S1**.

**Figure 5.**
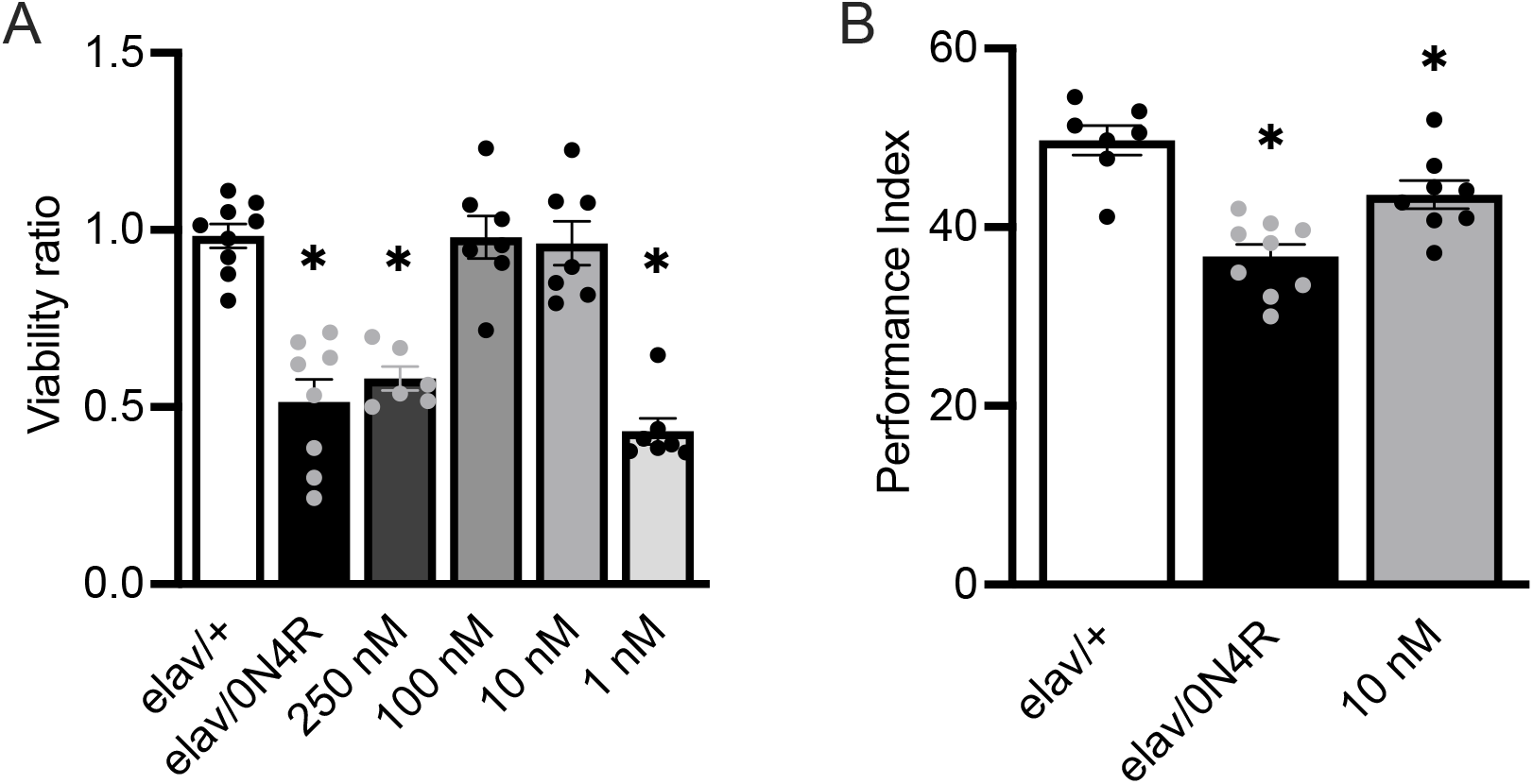
Effects of a polymorph-specific tau aggregation inhibitors (I1.21) on flies expressing human 0N4R tau. (**A**) Viability of flies expressing pan-neuronally human 0N4R tau (hTau0N4R) treated with vehicle (elav/0N4R) and increasing concentrations of compound I1.21. Control animals are driver heterozygotes and do not express the tau transgene (elav/+). (**B**) Twenty-four-hour spaced conditioning memory (PSD-M) performance of flies with (grey bar) or without (black bar) 10 nM of compound I1.21. For all the experiments bars indicate mean ± SEM and stars (*) indicate significant differences from both controls (elav/+). Statistical details are presented in Supplementary **Tables S1** and **S2**.

We then explored whether I1.21 treatment could impact tau-mediated neuronal dysfunction, particularly memory deficits in tau-expressing flies. As previously reported (*47*), pan-neuronal tau accumulation significantly decreases memory compared to controls (**Figure 5B**, ANOVA: F_(2.23)_=18.0, p=3×10^−5^; subsequent LSM: p=6×10^−6^ vs control). Treatment with I1.21 significantly improved the memory of tau-expressing flies relative to untreated tau flies (elav/0N4R) (**Figure 5B**, ANOVA:F_(2.23)_=18, p=3×10^−5^; subsequent LSM: p=3×10^−3^ vs elav/0N4R), although it did not fully restore it to the level of driver heterozygous controls (elav/+) (**Figure 5B**, ANOVA: F_(2.23)_=18, p=3×10^−5^; subsequent LSM: p=0.01 vs elav/+). Neither I1.51 nor I1.114 improved memory performance (**Figure S9Aii**,**Bii**). Detailed statistics can be found in **Table S2**. Thus, I1.21 not only mitigates tau-mediated toxicity but also alleviates tau-associated neuronal dysfunction.

## Discussion

The goal of this work was to develop a framework for a structure-based machine learning approach to target tau aggregation in a morphology-specific manner. We combined molecular docking simulations with *in vitro* kinetic analysis to identify compounds that could bind the amyloid fibril structures formed by tau in Alzheimer’s disease (*16*). We focused on targeting a hydrophobic pocket in the structure of the tau fibrils adjacent to PHF6 (^306^VQIVYK^312^), a known critical region in the aggregation cascade (*18*).

The computational methodology for pocket finding, subsequent virtual screening and machine learning iterative optimisation was previously established and shown to be effective for identifying and optimising aggregation inhibitors (*35*). This approach leverages a combination of multiple docking approaches and a QSAR optimisation module that consists of two regressors, a random forest fitted to the aggregation data, and a Gaussian process regressor fitted to the residuals of the first regressor, allowing utilization of the prediction uncertainty as part of the acquirement score (*35*). Tuning the uncertainty weighting allows a balancing between exploration of new chemical space vs exploitation of known chemical space. Both regressors used molecular embeddings derived from a pretrained junction tree variational autoencoder (*48*).

A limitation was a heuristic approach to pocket identification (*35*), based on a combination of pocket finding (*40*) and pocket solubility assessment (*41*). Pocket finding is a generally challenging task and would benefit from the consensus of different computational approaches such as those used here, and molecular dynamics and machine learning based methods. A second limitation is the relatively narrow diversity that can be achieved in the optimisation module due to searching within a library of similar molecules to the initial hits. Diversity could be increased via generative modelling (*49*), but this approach carries a risk of lower hit rates due to the low generalisability that accompanies training on relatively small datasets. This pipeline represents a compromise between ensuring potency of predicted molecules where resources for experimental testing are limited, ease of obtaining the molecules for testing, and the diversity of the predicted molecules.

On the experimental side, our approach builds upon previous work that demonstrated that a 3-repeat isoform of tau, K12, could reproducibly adopt two distinct structures when seeded with AD or PiD brain homogenates (*26*). Such an assay provides an on-target tau conformer and an off-target control for strain-specific kinetic analyses. We reported the reproducible propagation of two unique K12 conformers seeded by AD and PiD brain homogenates in the absence of heparin, a co-factor commonly required to promote tau aggregation *in vitro*. We showed through an array of approaches that the predicted inhibitors bind to tau fibrils in polymorph-specific assays. Fluorescence polarisation experiments confirmed the binding to the amyloid fibrils. PiD-seeded K12 reactions in the presence of AD-targeted small molecule inhibitors progressed at rates identical to control-spiked reactions, confirming conformer-specific binding.

The efficacy of the identified compounds in the *Drosophila* melanogaster tauopathy model highlights their potential as therapeutic candidates for targeting tau aggregation *in vivo*. In alignment with previous studies demonstrating the pivotal role of tau aggregation intermediates in driving toxicity (*50, 51*), treatment with I1.21 notably restored the viability and memory performance of tau-expressing flies to near-control levels. The observed differences in efficacy among the compounds may reflect variations in their bioavailability, stability, and metabolism within the *Drosophila* melanogaster model, with I1.21 potentially reaching effective concentrations in neuronal tissues more efficiently than I1.51 or I1.114. Furthermore, I1.21 was shown to inhibit both secondary nucleation and elongation processes, whereas I1.51 and I1.114 primarily impacted secondary nucleation. While the exact aggregation mechanism of human tau in *Drosophila* remains unclear (i.e. primary versus secondary nucleation), the ability of I1.21 to target multiple critical aggregation mechanisms, combined with its relatively high binding affinity, likely contributed to its superior in vivo efficacy in mitigating both tau toxicity and neuronal dysfunction in the *Drosophila* model. By disrupting aggregation pathways, I1.21 provides a promising framework for developing polymorph-specific inhibitors that may address the heterogeneity of tau pathology across neurodegenerative diseases.

## Conclusions

Tau aggregation is a major target for disease-modifying AD therapies (*8, 10*). Since tau self-assembles into a range of distinct amyloid fibril polymorphs underlying diverse neurodegenerative diseases (*16, 17, 19*), the study of aggregation *in vitro* should be concerned with inhibiting specific tau amyloid polymorphs (*19, 23*). Here we demonstrated that a kinetic analysis of AD tau-seeded aggregation could enhance the potency of small molecule aggregation inhibitors initially identified through docking to an exposed hydrophobic motif on the AD tau amyloid fibril. Iterative rounds of machine learning yielded a greater proportion of aggregation inhibitors compared to accelerators, strengthened potency, and produced common motifs among the most potent inhibitors. The best candidate compounds were demonstrated to specifically inhibit both secondary aggregation processes as well as elongation in polymorph-specific aggregation assays. Finally, when tested in an AD *Drosophila* model, the most potent compound rescued memory deficits and viability to the level of control flies. We anticipate that this approach will serve to identify further small molecules that can inhibit specific strains of tau and other amyloid-forming proteins.

## Methods

### Neuropathology and compliance with ethical standards

Procurement and neuropathology of brain samples utilised in this work was performed by Bernardino Ghetti, Indiana University School of Medicine. Brain samples in this work were obtained from deceased and de-identified consenting patients, requiring no further ethical disclosure. Briefly, one half of a patient brain was formalin fixed and the other half frozen. Diagnoses of fixed tissues were made using previously described immunohistochemical stains (*52*). Tissue samples in this study were collected from frontal cortex. 10% w/v brain homogenates of frontal cortex tissue were prepared in ice-cold PBS using 1 mm silica beads (BioSpec, 11079110z) and Beadbeater (BioSpec) or BeadMill 24 (Fischer). Homogenised samples were stored at −80 °C prior to thawing at room temperature for assay use. All Sporadic AD (sAD) and PiD brains listed in Table 1 in Ref. (*26*) were utilised during the optimisation stage of this work. Following the same sample numbering, data published in the main figures in this work include sAD2, PiD7, and CVD1 (*26*).

### Protein purification

K12 (Sequence: MGSSHHHHHHSSGLVPRGSHMQTAPVPMP DLKNVKSKIGSTENLKHQPGGGKVQIVYKPVDLSKVTSKAGSLGNIHHKPGGGQVEVKSEKLDFKDRVQSKIGSLDNITHVPGGGNKKIETHKLTFRENAKAKTDHGAEIVYKSPVVS) was purified as described previously (*26*). Briefly, the sequence for K12 tau with cysteine to serine mutation was cloned into a PET-28a vector and transformed into BL21(DE3) *E. coli*. Cells were grown and protein expression induced using an overnight autoinduction method described previously (*53*). Crude lysate was prepared as described previously with the addition of a boiling step prior to application to carboxylmethyl fast flow (CMFF) capture. K12 was eluted from CMFF resin over a 20 column volumes (CV) linear gradient from 100 – 500 mM NaCl. Pooled CMFF eluate was added to Sepharose High Performance (SPHP) resin and eluted over 40 CV linear gradient from 100-600 mM NaCl. SPHP fractions were pooled, precipitated in acetone, and dissolved in 8M GuHCl prior to size-exclusion chromatography (SEC) separation on a 26 × 600 mm Superdex 75 column equilibrated in 20 mM sodium phosphate, pH 7.4. The protein was lyophilised and frozen at −80 °C until use.

### First-generation seed amplification

Generation of AD-derived and PiD-derived K12 seeds was conducted as described previously with several modifications. Heparin was avoided in this study despite being previously required (*26*). NaF was replaced with 250 mM Na_3_Citrate; we previously showed heparin-free amplification of tau strains from brain homogenates with the use of Na_3_Citrate (*27, 54*). First-generation reactions were seeded with 1×10^−5^ concentration of brain homogenates in the presence of 4 μM K12, 10 μM ThT, 250 mM Na_3_citrate, 10 mM HEPES, pH 7.4. Reactions were subjected to rounds of 60 s shaking (500 rpm, orbital) and 60 s rest with periodic ThT readings every 15 min at 37 °C in a 384-well Nunc microplate (non-treated polymer base #242764) in a BMG FluoStar lite with aluminium sealing cover to prevent evaporation. Fibrils were harvested by scraping and pooling reaction contents once ThT fluorescence reached plateau > 20 h.

### Kinetic analysis of AD-derived and PiD-derived aggregation assays

One aliquot of purified and lyophilised K12 were dissolved in 1 mL 6M GuHCl prior to SEC separation on Superdex 75 10/300 column equilibrated in 20 mM sodium phosphate, pH 7.4. The protein was collected and diluted to obtain a series of concentrations ranging from 15.0 to 3.2 μM in 10 μM ThT, 250 mM Na_3_citrate, 10 mM HEPES buffered at pH 7.4. The initial kinetic model of AD-derived tau polymorph was created by reacting a concentration series of monomeric K12 tau with 1% preformed first-generation K12 tau fibrils initially seeded with AD brain homogenates. Kinetic analysis was conducted in quiescent conditions, with only mild agitation imposed by the moving of the plate in the plate reader (reading cycle = 15 min, with 1 min reads ad 1 min rest) to avoid biasing models with excessive fragmentation induced by shaking and shearing of fibrils. Half-times reported in **Figure 4** represent time to half max fluorescence intensity determined by simple sigmoidal dose-response curves fitted in GraphPad PRISM. The fitting of a power function to the half-time as a function of concentration yielded the exponent γ. Initial saturating secondary models were supported by γ values of −0.5 at lower monomeric concentrations of K12, following the AmyloFit protocol (*32*). Simpler models were chosen for initial analysis to avoid over-fitting of data. Addition of molar-equivalents (1X, 2X, 4X) of compounds to AD-derived kinetic reactions (1% DMSO) allowed for observation of kinetic modifications of reaction cascades in the presence of small molecules.

### ATR-FTIR analysis

Fibrils recovered from K12 first-generation amplification reactions were centrifuged for 10 min at 21,000 g and exchanged into water prior to ATR-FTIR analysis to avoid spectral contribution from buffer components. A Bruker Vertex 70 FTIR with diamond ATR sample attachment was used. Scans were taken from 800 cm^-1^ to 4,000 cm^-1^ with 100 replicates, step 4 Hz, apodization strong, correction: Happ-Genzel. Spectra were normalized to amide I intensity, and second derivatives were taken with 9 points for slope analysis.

### Computational docking and iterative machine learning

First, we selected a binding site on the tau fibrils. To achieve this goal, we analysed a structure of a tau fibril (PDB ID: 503l) (*16*) using Fpocket (*40*) which identifies potential binding pockets based on volume criteria. We identified a pocket on the fibril surface (encompassing residues V313, Asp314, Leu315), which had high surface exposure, necessary for secondary nucleation and high hydrophobicity, as identified by CamSol (*41*) (**Figure S2**), allowing the pocket to participate in aggregation. For the selection of screening compounds, we used the ZINC library, which contains a set of over 230 million purchasable compounds for screening (*42*). To prioritize the chemical space of small molecules considered in the docking calculations, central nervous system multiparameter optimisation (CNS MPO) criteria (*43*) were applied, effectively reducing the space to ∼2 million compounds. In particular, CNS MPO has been shown to correlate with key in vitro attributes of drug discovery, and thus using this filter potentially enables the identification of compounds with better physicochemical and pharmacokinetic properties pertaining to brain penetration, where tau is localized. We further subjected these compounds to docking calculation against the binding site identified above using AutoDock Vina (*44*). To increase the confidence of the calculations, the top-scoring 10000 small molecules were selected and docked against the same tau binding site, using FRED (*45*) (OpenEye Scientific Software). The top-scoring, common 1000 compounds in both docking protocols were selected and clustered using Tanimoto clustering, leading to a list of 130. Molecules were then obtained and tested experimentally in aggregation and binding experiments, before application of the iterative machine learning procedure as outlined in the online repository https://github.com/rohorne07/Iterate.

### Preparation of the compounds

The centroids from the above 130 clusters were selected for experimental validation. Compounds were purchased from Molport (Riga, Latvia), and in the cases for which centroids were not available for purchase, the compounds in the clusters with the closest chemical structures were used as the representative compounds instead. In the end, a total of 102 compounds were purchased (centroids and alternative compounds in 28 clusters were all not available for purchase) and then prepared in DMSO to a stock of 5 mM. Stocks were diluted in DMSO to 100-fold above the final desired final concentration, before addition to aggregation reactions at 100-fold dilution (1% DMSO). All chemicals used were purchased at the highest purity available (>90% in purity).

### Fluorescence polarization

10 µM of each molecule was incubated with increasing concentrations of K12 fibrils in the same buffer as used for kinetic experiments, supplemented with 1% DMSO. After incubation, the samples were pipetted into a 96-well half-area, black/clear flat bottom polystyrene nonbinding surface (NBS) microplate (Corning 3881). The fluorescence polarisation of the molecule was monitored using a plate reader (CLARIOstar, BMG Labtech, Aylesbury, UK) under quiescent conditions at room temperature, using a 360 nm excitation filter and a 520 nm emission filter.

### Recombinant Aß42 expression

The recombinant Aß42 peptide (MDAEFRHDSGY EVHHQKLVFF AEDVGSNKGA IIGLMVGGVV IA), here called Aß42, was expressed in the *E. coli* BL21 Gold (DE3) strain (Stratagene, CA, U.S.A.) and purified as described previously (*55*). Briefly, the purification procedure involved sonication of *E. coli* cells, dissolution of inclusion bodies in 8 M urea, and ion exchange in batch mode on diethylaminoethyl cellulose resin followed by lyophylisation. The lyophilised fractions were further purified using Superdex 75 HR 26/60 column (GE Healthcare, Buckinghamshire, U.K.) and eluates were analysed using SDS-PAGE for the presence of the desired peptide product. The fractions containing the recombinant peptide were combined, frozen using liquid nitrogen, and lyophilised again.

### Aß42 aggregation kinetics

Solutions of monomeric Aß42 were prepared by dissolving the lyophilized Aß42 peptide in 6 M guanidinium hydrocholoride (GuHCl). Monomeric forms were purified from potential oligomeric species and salt using a Superdex 75 10/300 GL column (GE Healthcare) at a flowrate of 0.5 mL/min, and were eluted in 20 mM sodium phosphate 200 µM EDTA, 0.02% NaN_3_, pH 8. The centre of the peak was collected, and the peptide concentration was determined from the absorbance of the integrated peak area using ε280 = 1490 l mol^-1^ cm^-1^. The obtained monomer was diluted with buffer to the desired concentration and supplemented with 20 μM thioflavin T (ThT) from a 2 mM stock. Each sample was then pipetted into multiple wells of a 96-well half-area, low-binding, clear bottom and PEG coated plate (Corning 3881), 80 µL per well, in the absence and the presence of different molar-equivalents of small molecules in 1% DMSO or 1% DMSO alone as a negative control. Assays were initiated by placing the 96-well plate at 37 ºC under quiescent conditions in a plate reader (Fluostar Omega, Fluostar Optima or Fluostar Galaxy, BMGLabtech, Offenburg, Germany). The ThT fluorescence was measured through the bottom of the plate using a 440 nm excitation filter and a 480 nm emission filter.

### *Drosophila melanogaster* culture and strains

Drosophila crosses were carried out in bulk using standard wheat-flour-sugar food, supplemented with soy flour and CaCl_2_. Cultures were maintained at 25 °C, with 50-70% humidity, and a 12 h light/dark cycle unless otherwise specified. Adult-specific pan-neuronal transgene expression was induced using the ElavC^155^-GAL4 driver, as described previously. The UAS-htau0N4R (human tau 0N4R) fly lines wewre a gift from Dr. Stefan Thor (Linkoping University)(*36*) and from Dr. Mel Feany (Harvard Medical School)(*37*).

### Viability assays

Drosophila crosses were maintained on standard fly food with or without the specified concentrations of the three compounds. To determine their effect on the viability of tau flies, five transgenic tau females(*37*) were crossed with three ElavC^155^-GAL4 males (Elav is on the X chromosome). Simultaneously, five *w*^*1118*^ females were crossed with three ElavC^155^-GAL4 males to serve as a control for the female-to-male ratio of their progeny. After 24 h, the flies were transferred to new vials, allowed to lay eggs for three days, and then discarded. The number of females versus males was counted when adults emerged. Each assessment was conducted at least five times, with five females each.

### Behavioural analyses

Flies expressing UAS-hTau0N4R(*36*) under the control of ElavC^155^-Gal4 driver were raised at 25 °C on standard fly food, supplemented with or without 10 nM of the indicated compound. Untreated driver heterozygotes were used as controls. All progenies were separated into groups of 50–70 mixed-sex flies. Olfactory aversive conditioning was performed as previously described [2, 4], using benzaldehyde (BNZ) and 3-octanol (OCT) diluted in isopropyl myristate (6% v/v for BNZ and 50% v/v for OCT) as conditioned stimuli (CS1 and CS2) with 90 V electric shocks as unconditioned stimuli (US). One hour before training, flies were transferred to fresh food vials. To assess 24 h memory after Spaced Conditioning, flies underwent 12 US/CS pairings per round and five training cycles with a 15 min rest interval between cycles, then were kept at 18 °C for 24 h before testing. In all experiments, two groups of the same genotype were trained simultaneously with the CS1 and CS2 odors switched. Both groups were tested in a T-maze apparatus, allowing them to choose between the two odors for 90 seconds. A performance index (PI) was calculated as previously described (*46*) and represents n = 1.

### Experimental design and statistical analyses

All experiments were conducted with both control and experimental genotypes tested in the same session using a balanced design. The genotypes were trained and tested in a random order. Behavioural experiment performance indexes were analysed parametrically using JMP 7.1 statistical software (SAS) and plotted with GraphPad Prism 9.5 software. After an initial positive ANOVA, means were compared with the control using planned multiple comparisons through the least squares means (LSM). The means and SEMs of viability were compared to those of the designated control using Dunnett’s test. All statistical details are provided in the text and relevant tables.

## Acknowledgements

We extend a most grateful acknowledgment to Bernardino Ghetti at Indiana University for his continued support of this work by performing neuropathology and providing tissue specimens critical to this work. This work was supported by the UKRI (10059436, 10061100). We acknowledge the expertise of Dr. Heather Greer in the microscopy core of the Department of Chemistry. Her helpful discussions on negative stain TEM imaging as well as acquisition of beautiful micrographs strengthened this manuscript. We would also like to thank ARCHER, MARCOPOLO and CIRCE high performance computing resources for the computer time. ZFB would like to acknowledge the Federation of European Biochemical Societies (FEBS) for financial support (LTF). MM was funded by the NIH/Cambridge scholars’ program.

## Notes

### Competing Interest Statement

The authors have declared no competing interest.

### Summary of Updates

One of the authors was not included in the online title, although present in the PDF.

## References

1. H. Braak, E. Braak, Neuropathological stageing of Alzheimer-related changes. Acta Neuropathol. 82, 239–259 (1991).

2. D. J. Selkoe, J. Hardy, The amyloid hypothesis of Alzheimer’s disease at 25 years. EMBO Mol. Med. 8, 595–608 (2016).

3. H. Hampel, J. Hardy, K. Blennow, C. Chen, G. Perry, S. H. Kim, V. L. Villemagne, P. Aisen, M. Vendruscolo, T. Iwatsubo, The amyloid-β pathway in Alzheimer’s disease. Mol. Psychiatry 26, 5481–5503 (2021).

4. C. R. Jack Jr, D. A. Bennett, K. Blennow, M. C. Carrillo, B. Dunn, S. B. Haeberlein, D. M. Holtzman, W. Jagust, F. Jessen, J. Karlawish, NIA‐AA research framework: toward a biological definition of Alzheimer’s disease. Alzheimers Dement. 14, 535–562 (2018).

5. L. C. Walker, M. I. Diamond, K.E. Duff, B. T. Hyman, Mechanisms of protein seeding in neurodegenerative diseases. JAMA Neurol. 70, 304–310 (2013).

6. J. Brettschneider, K. D. Tredici, V. M.-Y. Lee, J. Q. Trojanowski, Spreading of pathology in neurodegenerative diseases: a focus on human studies. Nat. Rev. Neurosci. 16, 109–120 (2015).

7. S. Wegmann, R. E. Bennett, L. Delorme, A. B. Robbins, M. Hu, D. MacKenzie, M. J. Kirk, J. Schiantarelli, N. Tunio, A. C. Amaral, Experimental evidence for the age dependence of tau protein spread in the brain. Sci. Adv. 5, eaaw6404 (2019).

8. J. Cummings, Y. Zhou, G. Lee, K. Zhong, J. Fonseca, F. Cheng, Alzheimer’s disease drug development pipeline: 2024. Alzheimers Dement. 10, e12465 (2024).

9. C. H. Van Dyck, C. J. Swanson, P. Aisen, R. J. Bateman, C. Chen, M. Gee, M. Kanekiyo, D. Li, L. Reyderman, S. Cohen, Lecanemab in early Alzheimer’s disease. N. Engl. J. Med. 388, 9–21 (2023).

10. E. E. Congdon, C. Ji, A. M. Tetlow, Y. Jiang, E. M. Sigurdsson, Tau-targeting therapies for Alzheimer disease: current status and future directions. Nat. Rev. Neurol. 19, 715–736 (2023).

11. S. I. Cohen, S. Linse, L. M. Luheshi, E. Hellstrand, D. A. White, L. Rajah, D. E. Otzen, M. Vendruscolo, C. M. Dobson, T. P. Knowles, Proliferation of amyloid-β42 aggregates occurs through a secondary nucleation mechanism. Proc. Natl. Acad. Sci. USA 110, 9758–9763 (2013).

12. T. P. Knowles, M. Vendruscolo, C. M. Dobson, The amyloid state and its association with protein misfolding diseases. Nat. Rev. Mol. Cell Biol. 15, 384–396 (2014).

13. D. Thacker, K. Sanagavarapu, B. Frohm, G. Meisl, T. P. Knowles, S. Linse, The role of fibril structure and surface hydrophobicity in secondary nucleation of amyloid fibrils. Proc. Natl. Acad. Sci. USA 117, 25272–25283 (2020).

14. M. Törnquist, T. C. Michaels, K. Sanagavarapu, X. Yang, G. Meisl, S. I. Cohen, T. P. Knowles, S. Linse, Secondary nucleation in amyloid formation. Chem. Comm. 54, 8667–8684 (2018).

15. K. Sanagavarapu, G. Meisl, V. Lattanzi, K. Bernfur, B. Frohm, U. Olsson, T. P. Knowles, A. Malmendal, S. Linse, Serine phosphorylation mimics of Aβ form distinct, non-cross-seeding fibril morphs. Chem. Sci. 15, 19142–19159 (2024).

16. A. W. Fitzpatrick, B. Falcon, S. He, A. G. Murzin, G. Murshudov, H. J. Garringer, R. A. Crowther, B. Ghetti, M. Goedert, S. H. Scheres, Cryo-EM structures of tau filaments from Alzheimer’s disease. Nature 547, 185–190 (2017).

17. Y. Shi, W. Zhang, Y. Yang, A. G. Murzin, B. Falcon, A. Kotecha, M. van Beers, A. Tarutani, F. Kametani, H. J. Garringer, Structure-based classification of tauopathies. Nature 598, 359–363 (2021).

18. S. Lövestam, D. Li, J.L. Wagstaff, A. Kotecha, D. Kimanius, S. H. McLaughlin, A. G. Murzin, S. M. Freund, M. Goedert, S. H. Scheres, Disease-specific tau filaments assemble via polymorphic intermediates. Nature 625, 119–125 (2024).

19. S. Lövestam, F. A. Koh, B. van Knippenberg, A. Kotecha, A. G. Murzin, M. Goedert, S. H. Scheres, Assembly of recombinant tau into filaments identical to those of Alzheimer’s disease and chronic traumatic encephalopathy. eLife 11, e76494 (2022).

20. W. Zhang, B. Falcon, A. G. Murzin, J. Fan, R. A. Crowther, M. Goedert, S. H. Scheres, Heparin-induced tau filaments are polymorphic and di_er from those in Alzheimer’s and Pick’s diseases. eLife 8, e43584 (2019).

21. P. Chakraborty, G. Rivière, S. Liu, A. I. de Opakua, R. Dervişoğlu, A. Hebestreit, L. B. Andreas, I. M. Vorberg, M. Zweckstetter, Co-factor-free aggregation of tau into seeding-competent RNA-sequestering amyloid fibrils. Nat. Comm. 12, 4231 (2021).

22. P. Duan, A. J. Dregni, H. Xu, L. Changolkar, V. M. Lee, E. B. Lee, M. Hong, Alzheimer’s disease seeded tau forms paired helical filaments yet lacks seeding potential. J. Biol. Chem. 300, 107730 (2024).

23. S. Lövestam, M. Schweighauser, T. Matsubara, S. Murayama, T. Tomita, T. Ando, K. Hasegawa, M. Yoshida, A. Tarutani, M. Hasegawa, Seeded assembly in vitro does not replicate the structures of α‐synuclein filaments from multiple system atrophy. FEBS Open Bio 11, 999–1013 (2021).

24. A. Tarutani, S. Lövestam, X. Zhang, A. Kotecha, A. C. Robinson, D. M. Mann, Y. Saito, S. Murayama, T. Tomita, M. Goedert, Cryo‐EM structures of tau filaments from SH‐SY5Y cells seeded with brain extracts from cases of Alzheimer’s disease and corticobasal degeneration. FEBS Open Bio 13, 1394–1404 (2023).

25. D. C. Rodriguez Camargo, E. Sileikis, S. Chia, E. Axell, K. Bernfur, R. L. Cataldi, S. I. Cohen, G. Meisl, J. Habchi, T. P. Knowles, Proliferation of tau 304–380 fragment aggregates through autocatalytic secondary nucleation. ACS Chem. Neurosci. 12, 4406–4415 (2021).

26. M. A. Metrick, N. d. C. Ferreira, E. Saijo, A. Kraus, K. Newell, G. Zanusso, M. Vendruscolo, B. Ghetti, B. Caughey, A single ultrasensitive assay for detection and discrimination of tau aggregates of Alzheimer and Pick diseases. Acta Neuropathol. Commun. 8, 1–13 (2020).

27. M. A. Metrick, N. do Carmo Ferreira, E. Saijo, A. G. Hughson, A. Kraus, C. Orrú, M. W. Miller, G. Zanusso, B. Ghetti, M. Vendruscolo, Million-fold sensitivity enhancement in proteopathic seed amplification assays for biospecimens by Hofmeister ion comparisons. Proc. Natl. Acad. Sci. USA 116, 23029–23039 (2019).

28. Y. Yang, Y. Shi, M. Schweighauser, X. Zhang, A. Kotecha, A. G. Murzin, H. J. Garringer, P. W. Cullinane, Y. Saito, T. Foroud, Structures of α-synuclein filaments from human brains with Lewy pathology. Nature 610, 791–795 (2022).

29. M. Schweighauser, Y. Shi, A. Tarutani, F. Kametani, A. G. Murzin, B. Ghetti, T. Matsubara, T. Tomita, T. Ando, K. Hasegawa, Structures of α-synuclein filaments from multiple system atrophy. Nature 585, 464–469 (2020).

30. E. Axell, J. Hu, M. Lindberg, A. J. Dear, L. Ortigosa-Pascual, E. A. Andrzejewska, G. Šneiderienė, D. Thacker, T. P. Knowles, E. Sparr, The role of shear forces in primary and secondary nucleation of amyloid fibrils. Proc. Natl. Acad. Sci. USA 121, e2322572121 (2024).

31. T. P. Knowles, C. A. Waudby, G. L. Devlin, S. I. Cohen, A. Aguzzi, M. Vendruscolo, E. M. Terentjev, M. E. Welland, C. M. Dobson, An analytical solution to the kinetics of breakable filament assembly. Science 326, 1533–1537 (2009).

32. G. Meisl, J. B. Kirkegaard, P. Arosio, T. C. Michaels, M. Vendruscolo, C. M. Dobson, S. Linse, T. P. Knowles, Molecular mechanisms of protein aggregation from global fitting of kinetic models. Nat. Protoc. 11, 252–272 (2016).

33. S. Chia, J. Habchi, T. C. Michaels, S. I. Cohen, S. Linse, C. M. Dobson, T. P. Knowles, M. Vendruscolo, SAR by kinetics for drug discovery in protein misfolding diseases. Proc. Natl. Acad. Sci. USA 115, 10245–10250 (2018).

34. J. Habchi, S. Chia, R. Limbocker, B. Mannini, M. Ahn, M. Perni, O. Hansson, P. Arosio, J. R. Kumita, P. K. Challa, Systematic development of small molecules to inhibit specific microscopic steps of Aβ42 aggregation in Alzheimer’s disease. Proc. Natl. Acad. Sci. USA 114, E200–E208 (2017).

35. R. I. Horne, E. A. Andrzejewska, P. Alam, Z. F. Brotzakis, A. Srivastava, A. Aubert, M. Nowinska, R. C. Gregory, R. Staats, A. Possenti, Discovery of potent inhibitors of α-synuclein aggregation using structure-based iterative learning. Nat. Chem. Biol. 20, 634–645 (2024).

36. J. Fernius, A. Starkenberg, M. Pokrzywa, S. Thor, Human TTBK1, TTBK2 and MARK1 kinase toxicity in Drosophila melanogaster is exacerbated by co-expression of human Tau. Biol. Open 6, 1013–1023 (2017).

37. C. W. Wittmann, M. F. Wszolek, J. M. Shulman, P. M. Salvaterra, J. Lewis, M. Hutton, M. B. Feany, Tauopathy in Drosophila: neurodegeneration without neurofibrillary tangles. Science 293, 711–714 (2001).

38. S. Chia, Z. Faidon Brotzakis, R. I. Horne, A. Possenti, B. Mannini, R. Cataldi, M. Nowinska, R. Staats, S. Linse, T. P. Knowles, Structure-based discovery of small-molecule inhibitors of the autocatalytic proliferation of α-Synuclein aggregates. Mol. Pharmaceutics 20, 183–193 (2022).

39. R. Staats, Z. F. Brotzakis, S. Chia, R. I. Horne, M. Vendruscolo, Optimization of a small molecule inhibitor of secondary nucleation in α-synuclein aggregation. Front. Mol. Biosci. 10, 1155753 (2023).

40. V. Le Guilloux, P. Schmidtke, P. Tuffery, Fpocket: an open source platform for ligand pocket detection. BMC Bioinformatics 10, 1–11 (2009).

41. P. Sormanni, F. A. Aprile, M. Vendruscolo, The CamSol method of rational design of protein mutants with enhanced solubility. J. Mol. Biol. 427, 478–490 (2015).

42. J. J. Irwin, B. K. Shoichet, ZINC - a free database of commercially available compounds for virtual screening. J. Chem. Inf. Model. 45, 177–182 (2005).

43. T. T. Wager, X. Hou, P. R. Verhoest, A. Villalobos, Central nervous system multiparameter optimization desirability: application in drug discovery. ACS Chem. Neurosci. 7, 767–775 (2016).

44. O. Trott, A. J. Olson, AutoDock Vina: improving the speed and accuracy of docking with a new scoring function, e_icient optimization, and multithreading. J. Comp. Chem. 31, 455–461 (2010).

45. M. McGann, FRED pose prediction and virtual screening accuracy. J. Chem. Inf. Model. 51, 578–596 (2011).

46. E. Prifti, E. N. Tsakiri, E. Vourkou, G. Stamatakis, M. Samiotaki, E. M. Skoulakis, K. Papanikolopoulou, Mical modulates Tau toxicity via cysteine oxidation in vivo. Acta Neuropathol. Commun. 10, 44 (2022).

47. E. Vourkou, V. Paspaliaris, A. Bourouliti, M.-C. Zerva, E. Prifti, K. Papanikolopoulou, E. M. Skoulakis, Differential effects of human tau isoforms to neuronal dysfunction and toxicity in the drosophila CNS. Int. J. Mol. Sci. 23, 12985 (2022).

48. W. Jin, R. Barzilay, T. Jaakkola, Junction tree variational autoencoder for molecular graph generation. International conference on machine learning, 2323–2332 (2018).

49. B. A. Koscher, R. B. Canty, M. A. McDonald, K. P. Greenman, C. J. McGill, C. L. Bilodeau, W. Jin, H. Wu, F. H. Vermeire, B. Jin, Autonomous, multiproperty-driven molecular discovery: From predictions to measurements and back. Science 382, eadi1407 (2023).

50. C. Ballatore, V. M.-Y. Lee, J. Q. Trojanowski, Tau-mediated neurodegeneration in Alzheimer’s disease and related disorders. Nat. Rev. Neurosci. 8, 663–672 (2007).

51. E. Vourkou, E. D. R. Ortega, S. Mahajan, A. Mudher, E. M. Skoulakis, Human Tau Aggregates Are Permissive to Protein Synthesis-Dependent Memory in Drosophila Tauopathy Models. J. Neurosci. 43, 2988–3006 (2023).

52. C. Bouras, P. R. Hof, J. H. Morrison, Neurofibrillary tangle densities in the hippocampal formation in a non-demented population define subgroups of patients with differential early pathologic changes. Neurosci. Lett. 153, 131–135 (1993).

53. F. W. Studier, Protein production by auto-induction in high-density shaking cultures. Protein Expr. Purif. 41, 207–234 (2005).

54. S. Koga, M. A. Metrick, L. I. Golbe, A. Santambrogio, M. Kim, A. I. Soto-Beasley, R. L. Walton, M. C. Baker, C. F. De Castro, M. DeTure, Case report of a patient with unclassified tauopathy with molecular and neuropathological features of both progressive supranuclear palsy and corticobasal degeneration. Acta Neuropathol. Commun. 11, 88 (2023).

55. S. Linse, Expression and purification of intrinsically disordered Aβ peptide and setup of reproducible aggregation kinetics experiment. Methods Mol Biol. 2141, 731–754 (2020).

