## Supplementary Information for "Design of Tau Aggregation Inhibitors Using Iterative Machine Learning and a Polymorph-Specific Brain-Seeded Fibril Amplification Assay"

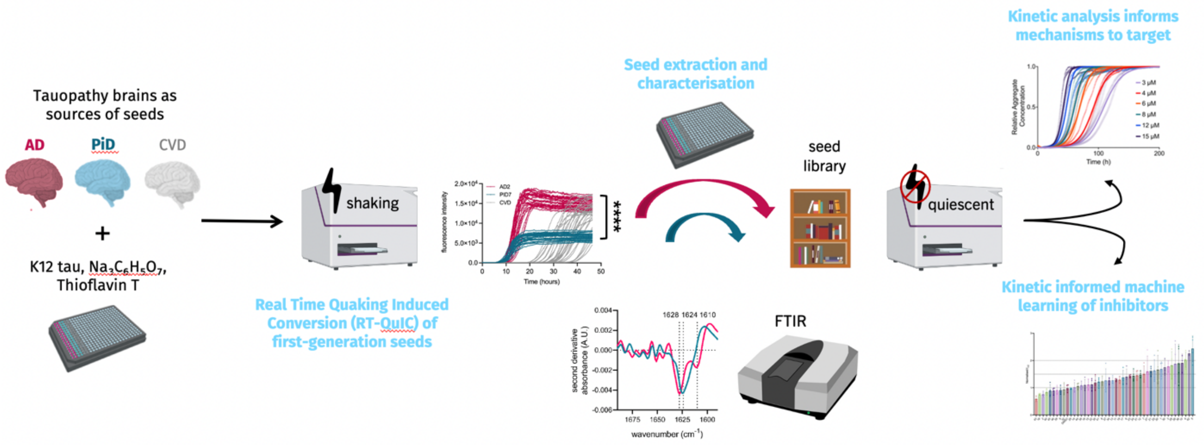


**Figure S1**. **Schematic illustration of the protocol for polymorph-specific drug design assay used in this work**. Brain-derived seeds are obtained from tauopathy and control brain homogenates and used to promote the *in vitro* conversion of recombinant K12 tau into aggregates. Recombinant seeds are then extracted from this aggregation reaction to form libraries for downstream biophysical characterization and use in drug design assays. In these assays, recombinant tau monomers are reacted with recombinant seed libraries in the presence of docked compounds over rounds of iterative machine learning, informed by kinetic analysis of inhibition. This approach yields potent polymorph-specific tau aggregation inhibitors. (AD, Alzheimer's disease; PiD, Pick's disease; CVD, cerebrovascular disease).

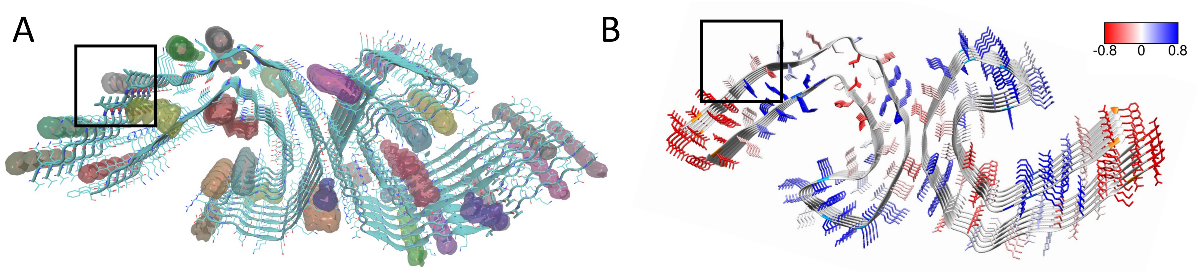


**Figure S2. Binding site predictions based on pocket identification and solubility analysis. (A)** Cavity-based binding pocket prediction using Fpocket (*1*). **(B)** Solubility-based scoring of the predicted binding pockets using CamSol (*2*). The black box indicates residues 313-315, where both solubility is low and cavity propensity is high. This approach was recently validated for the structure-based discovery of α-synuclein aggregation inhibitors (*3*).

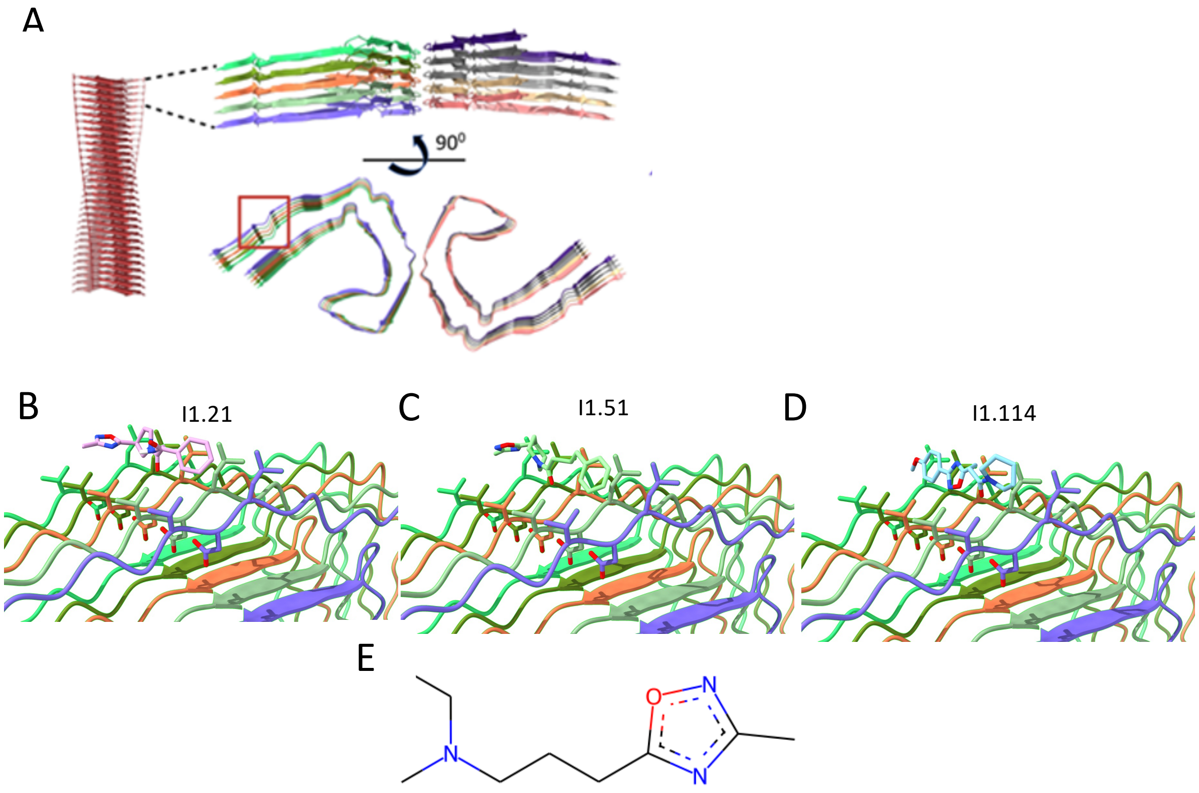


**Figure S3**. **Small molecule features**. (**A**) Schematic of the proposed binding mechanism, showing the cross section of the *ex vivo* AD tau fibril structure (*4*) and the targeted binding site. Binding of I1.21 in this pocket as predicted by AutoDock Vina is also shown. (**B-D**) Best docking poses for the three compounds (I1.21 (B), I1.51 (C), and I1.114 (C)) against the AD tau fibril (PDB:5O3L) at the binding site (V313-L315). (**E**) Maximum common substructure between compounds, determined by RDKit.

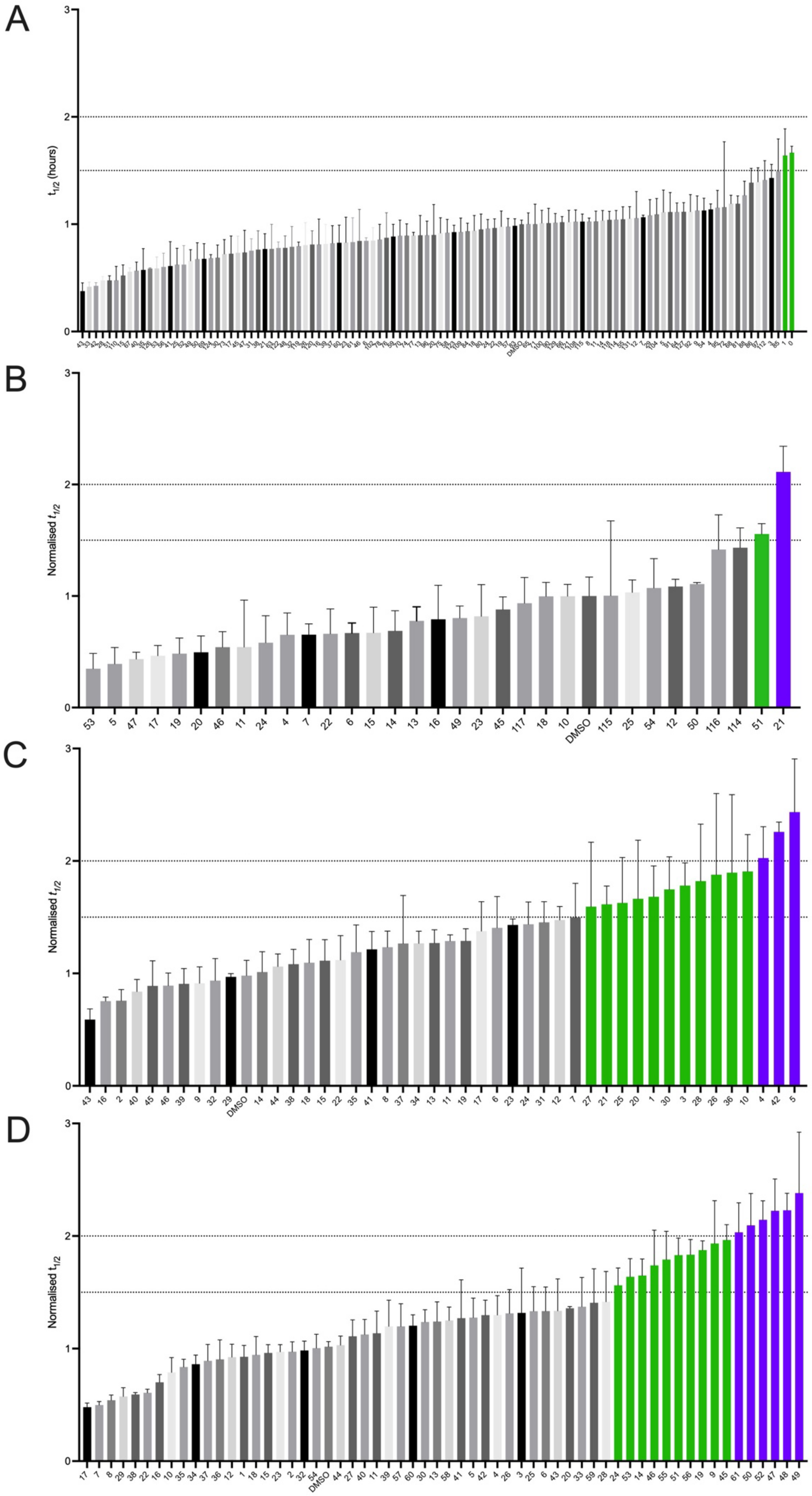
A
$t_{1/2}$ (hours)
B
Normalised $t_{1/2}$
53 5 47 17 19 20 46 11 24 4 7 22 6 15 14 13 16 49 23 45 117 18 10 DMSO 115 25 54 12 50 116 114 51 21
C
Normalised $t_{1/2}$
43 16 2 40 45 46 39 9 32 29 DMSO 14 44 38 18 15 22 35 41 8 37 34 13 11 19 17 6 23 24 31 12 7 27 21 25 20 1 30 3 28 26 36 10 4 42 5
D
Normalised $t_{1/2}$
17 7 8 29 38 22 16 10 35 34 37 36 12 1 18 15 23 2 32 54 DMSO 44 27 40 11 39 57 60 30 13 58 41 5 42 4 26 3 25 6 43 20 33 59 28 24 53 14 46 55 51 56 19 9 45 61 50 52 47 48 49

**Figure S4. Summary of the aggregation data for the compounds tested in this work in the presence of first-generation fibrils.** Bars represent the average half time ($t_{1/2}$) of 4 replicate reactions of 5 µM K12 tau monomer, 50 nM first-generation AD-seeded K12 fibrils in the presence of 1% DMSO (error=SD) and 20 µM molecules I1.21, I1.51 and I1.114; averages are normalised to DMSO = 1. Green bars represent reactions with $t_{1/2}$ > 1.5 greater or more than DMSO control, indicating aggregation inhibitors. Purple bars represent reactions with greater than two-fold $t_{1/2}$ prolongation. **(A)** $t_{1/2}$ values of AD-seeded K12 tau in the presence of original docked compounds. **(B)** $t_{1/2}$ values of the same reactions in the presence of compounds from first iteration of kinetic-informed machine learning. **(C)** As above for iteration two. **(D)** As above for iteration three.

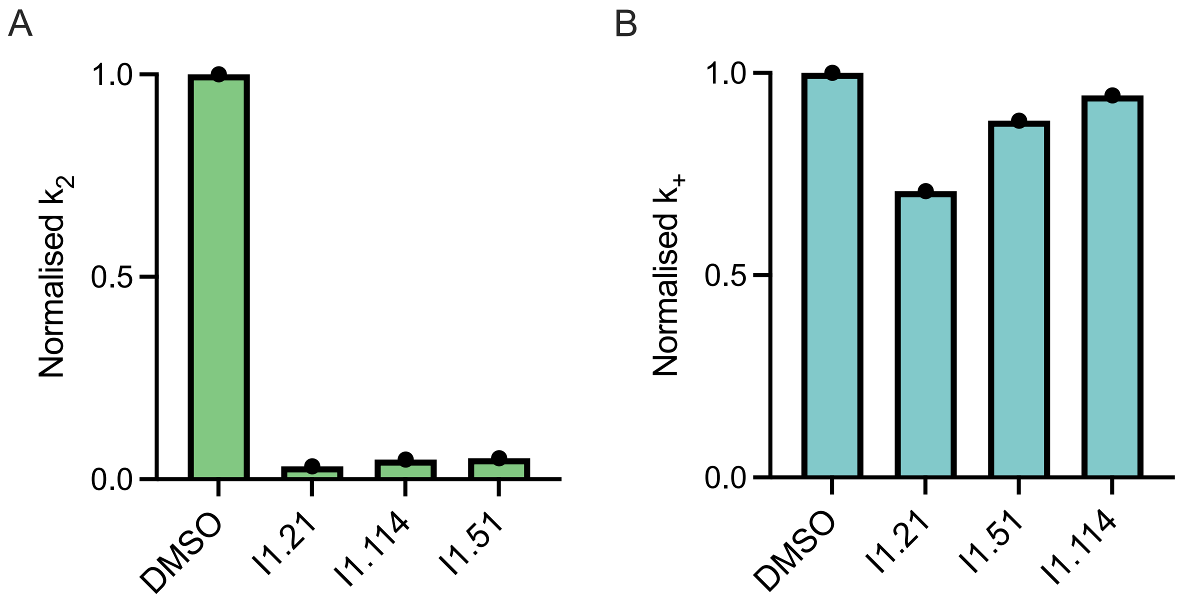


**Figure S5. Secondary nucleation rate constant ($k_2$) and elongation rate constat ($k_+$) for the molecules tested in this work.** $k_2$ (**A**, green) and $k_+$ (**B**, teal) values obtained when fitted aggregation curves in **Figure 4B** to a secondary nucleation model at 20 µM (4:1 stoichiometry) of I1.21, I1.51 and I1.114.

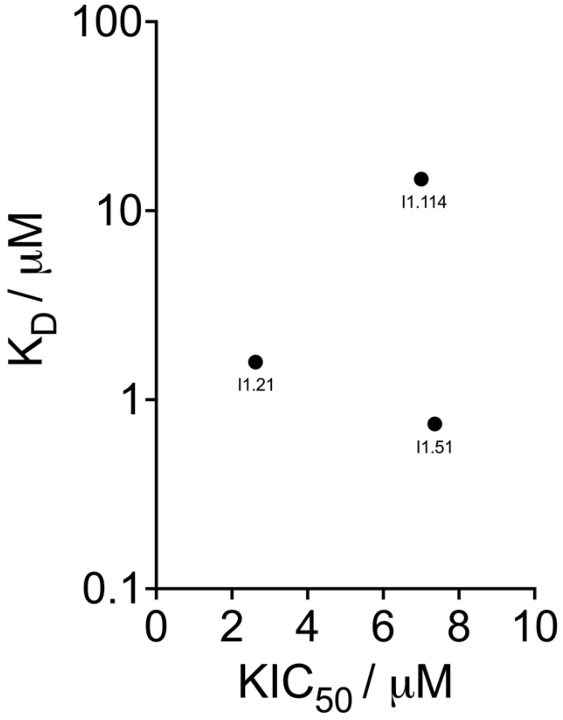


**Figure S6. Comparison of the $KIC_{50}$ and $K_D$ values for the 3 small molecules reported in this work.** The $K_D$ values report on the binding affinity of the small molecules for the tau fibrils, while the $KIC_{50}$ values quantify their potency as inhibitors of fibril proliferation.

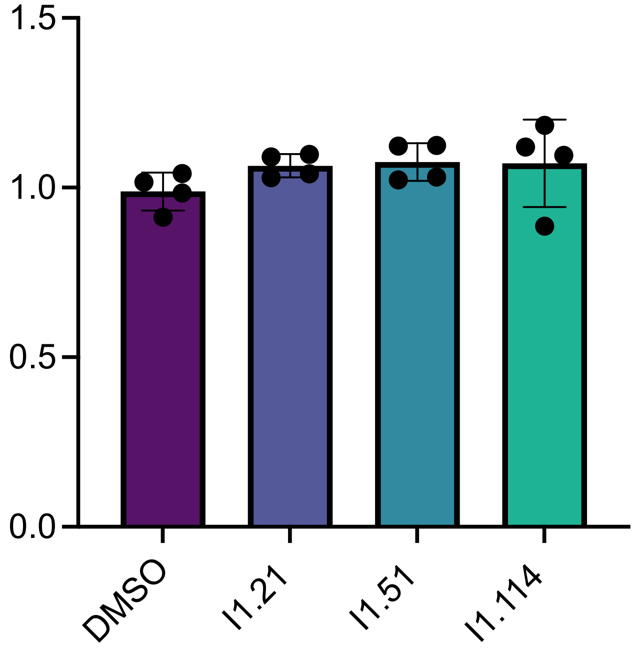


**Figure S7. PiD brain homongenate-seeded K12 aggregtion $t_{1/2}$ in the presence of the 3 small molecules reported in this work.** Normalised half-time ($t_{1/2}$) of PiD-seeded K12 reactions for the most potent AD-seeded K12 aggregation inhibitors described in this work.

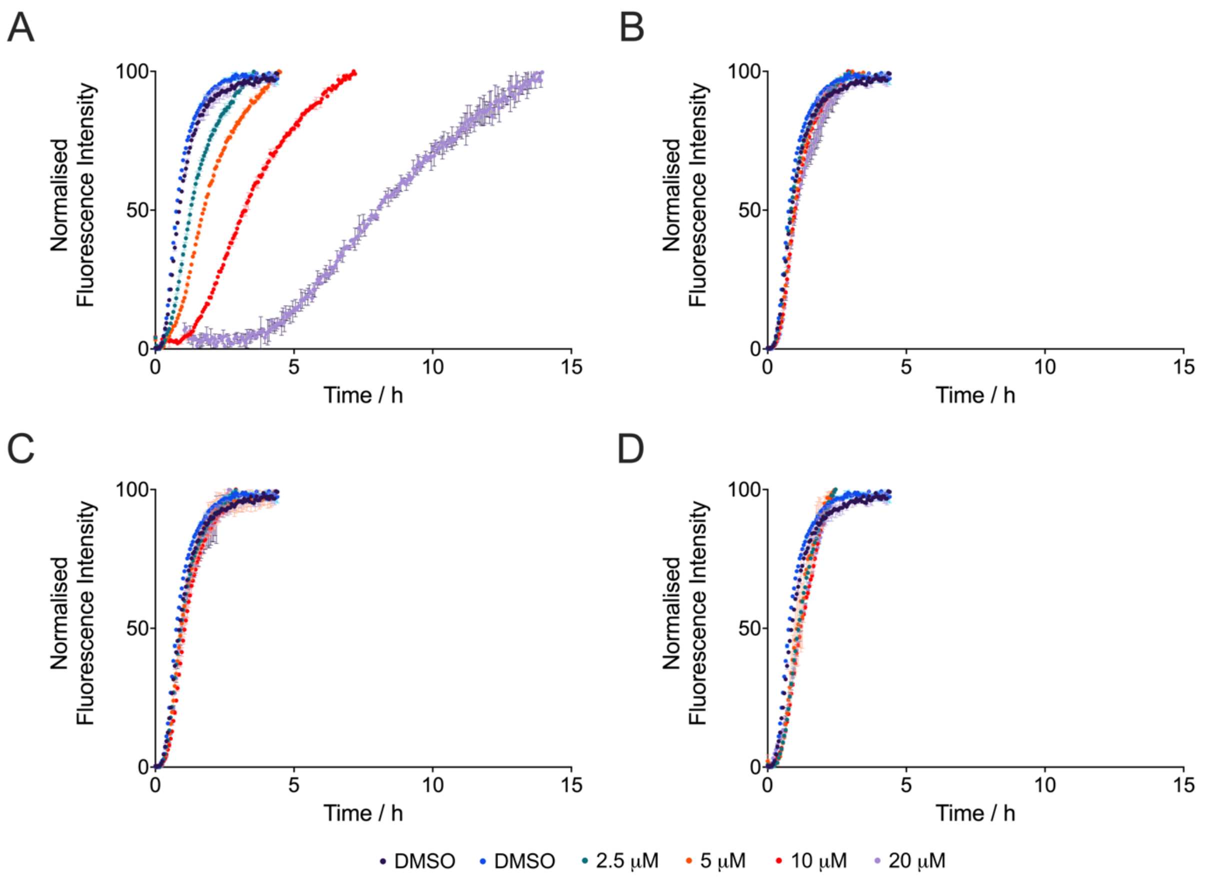


**Figure S8. Kinetic traces of Aß42 aggregation in the presence of the 3 small molecules reported in this work**. Aß42 (40 nM seed, 2 μM monomer) was aggregated in the presence of 1% DMSO (purple and blue points), and molecules at 2.5 μM (teal), 5 μM (orange), 10 μM (red), and 20 μM (lilac). (**A**) Adapalene, a positive control with previously reported anti-aggregation potency (*5*). (**B**) Compound I1.21. (**C**) Compound I1.51. (**D**) Compoud I1.114.

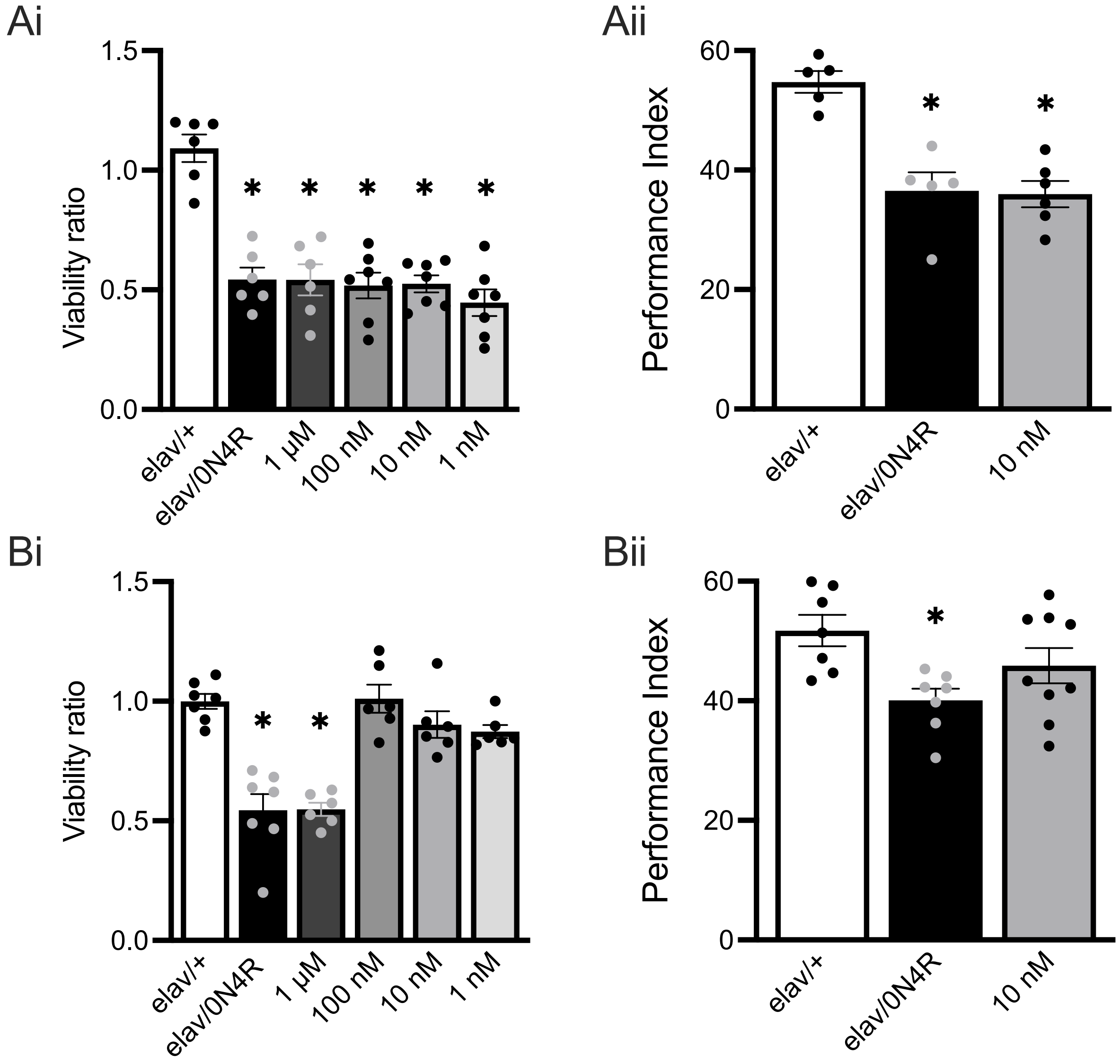


**Figure S9. Effects of two polymorph-specific tau aggregation inhibitors (I1.51 and I.114) on flies expressing human 0N4R tau**. **(Ai)** Viability of flies treated with vehicle (control) and increasing concentrations of compound I1.51. (**Aii**) Twenty-four-hour Spaced Conditioning memory (PSD-M) performance of flies with or without 10 μM of compound I1.51. **(Bi)** As above for I1.114. **(Bii)** As above for I1.114. For all the experiments bars indicate mean ± SEM and and stars (*) indicate significant differences from both controls (elav/+). Statistical details are presented in Supplementary **Table S1** and **S2**.

**Table S1: Comparisons to control elav$^{C155}$-Gal4> w$^{1118}$ using Dunnett's test.**

| **d I1.21** | **p-value** | **I1.51** | **p-value** | **I1.114** | **p-value** |
|---|---|---|---|---|---|
| elav/+ | 1 | elav/+ | 1 | elav/+ | 1 |
| elav/0N4R | 1e-07 | elav/0N4R | 2e-07 | elav/0N4R | 3e07 |
| tau 100 nM | 1 | tau 1 μM | 2e-07 | tau 100 nM | 1 |
| tau 10 nM | 1 | tau 10 nM | 4e-08 | tau 10 nM | 0.5 |
| tau 250 nM | 1e-05 | tau 100 nM | 3e-08 | tau 1 nM | 0.2 |
| tau 1 nM | 2e-09 | tau 1 nM | 0 | tau 1 μM | 7e-07 |

**Table S2: Comparisons to control elav$^{C155}$-Gal4> w$^{1118}$ and elav$^{C155}$-Gal4> 0N4R using LSM.**

| Genotype | Mean | ± SEM | p-value | F-ratio |
|---|---|---|---|---|
| **LTM 10nM d21** | | **ANOVA $F_{(2.23)}$=18 p=0.00003** | | |
| elav$^{C155}$-Gal4> w$^{1118}$ | 50 | ±2 | | |
| elav$^{C155}$-Gal4 >0N4R | 37 | ±1 | **6e-06** | 36 |
| elav$^{C155}$-Gal4 >0N4R I1.21 | 44 | ±2 | **0.01** | 7 |
| elav$^{C155}$-Gal4> 0N4R | 37 | ±1 | | |
| elav$^{C155}$-Gal4 >0N4R I1.21 | 44 | ±2 | **0.003** | 11 |
| **LTM 10nM d51** | | **ANOVA $F_{(2.15)}$=18 p=0.0002** | | |
| elav$^{C155}$-Gal4> w$^{1118}$ | 56 | ±3 | | |
| elav$^{C155}$-Gal4 >0N4R | 37 | ±3 | **0.0002** | 25 |
| elav$^{C155}$-Gal4 >0N4R I1.51 | 36 | ±2 | **0.0001** | 29 |
| elav$^{C155}$-Gal4> 0N4R | 37 | ±3 | | |
| elav$^{C155}$-Gal4 >0N4R I1.51 | 36 | ±2 | 0.9 | 0.02 |
| **LTM 10nM d114** | | **ANOVA $F_{(2.22)}$=4 p=0.03** | | |
| elav$^{C155}$-Gal4> w$^{1118}$ | 52 | ±3 | | |
| elav$^{C155}$-Gal4 >0N4R | 40 | ±2 | **0.008** | 9 |
| elav$^{C155}$-Gal4 >0N4R I1.114 | 46 | ±3 | 0.1 | 3 |
| elav$^{C155}$-Gal4> 0N4R | 40 | ±2 | | |
| elav$^{C155}$-Gal4 >0N4R I1.114 | 46 | ±3 | 0.1 | 2 |

### Supplementary References


1. V. Le Guilloux, P. Schmidtke, P. Tuffery, Fpocket: an open source platform for ligand pocket detection. *BMC Bioinformatics* **10**, 1-11 (2009).
2. P. Sormanni, F. A. Aprile, M. Vendruscolo, The CamSol method of rational design of protein mutants with enhanced solubility. *J. Mol. Biol.* **427**, 478-490 (2015).
3. S. Lövestam, D. Li, J. L. Wagstaff, A. Kotecha, D. Kimanius, S. H. McLaughlin, A. G. Murzin, S. M. Freund, M. Goedert, S. H. Scheres, Disease-specific tau filaments assemble via polymorphic intermediates. *Nature* **625**, 119-125 (2024).
4. A. W. Fitzpatrick, B. Falcon, S. He, A. G. Murzin, G. Murshudov, H. J. Garringer, R. A. Crowther, B. Ghetti, M. Goedert, S. H. Scheres, Cryo-EM structures of tau filaments from Alzheimer’s disease. *Nature* **547**, 185-190 (2017).
5. J. Habchi, S. Chia, R. Limbocker, B. Mannini, M. Ahn, M. Perni, O. Hansson, P. Arosio, J. R. Kumita, P. K. Challa, Systematic development of small molecules to inhibit specific microscopic steps of Aβ42 aggregation in Alzheimer’s disease. *Proc. Natl. Acad. Sci. USA* **114**, E200-E208 (2017).
